# Lipotoxicity Induces Beta Cell Small Extracellular Vesicle-mediated β-cell Dysfunction

**DOI:** 10.1101/2024.07.19.604353

**Authors:** Abhishek Roy, Alexandra Hoff, Tracy K. Her, Gallage Ariyaratne, Kamalnath Sankaran Rajagopalan, Matthew R. Brown, Alondra Soto-González, Aleksey V. Matveyenko, Naureen Javeed

## Abstract

Chronically elevated circulating excess free fatty acids (*i.e.* lipotoxicity) is a pathological process implicated in several metabolic disorders, including obesity-driven Type 2 diabetes (T2D). Lipotoxicity exerts detrimental effects on pancreatic islet β-cells by reducing glucose-stimulated insulin secretion (GSIS), altering β-cell transcriptional identity, and promoting apoptosis. While β-cell-derived small extracellular vesicles (sEV) have been shown to contribute to β-cell failure in T2D, their specific role in lipotoxicity-mediated β-cell failure remains to be elucidated. In this work, we demonstrate that lipotoxicity enhances the release of sEVs from β-cells, which exhibit altered proteomic and lipidomic profiles. These lipotoxic sEV induce β-cell dysfunction in healthy mouse and human islets and trigger significant islet transcriptional changes, including the upregulation of genes associated with the TGFβ/Smad3 pathway, as noted by RNA sequencing. Importantly, pharmacological inhibition of the TGFβI/II receptor improved lipotoxic sEV-induced β-cell dysfunction, underscoring their involvement in activating the TGFβ/Smad3 pathway during this process. We have comprehensively characterized lipotoxic β-cell sEVs and implicated their role in inducing β-cell functional failure in T2D. These findings highlight potential avenues for therapeutic interventions targeting sEV-mediated pathways to preserve β-cell health in metabolic disorders.

**Article Highlights:**

- Diabetogenic lipotoxic conditions enhance β-cell sEV release and induce alterations in both protein and lipid content.
- Global islet transcriptional changes and alterations in β-cell function were noted upon exposure to lipotoxic sEV.
- Lipotoxic sEV were shown to activate the TGFβ/Smad3 pathway and blockade of this pathway improved β-cell function.

---

Type 2 diabetes (T2D) is a multifactorial disease characterized by peripheral insulin resistance, coupled with dysfunction and progressive loss of β-cell mass^1–3^. Although complex in its pathophysiology, obesity is a well-established risk factor in the etiology of T2D. Overnutrition or dysregulation in lipid metabolism results in the accumulation of excess free fatty acid (FFA) in plasma, which profoundly affects the function and cellular metabolism of peripheral tissues^4^. Pancreatic β-cells are vulnerable to the deleterious effects of FFAs. Chronic exposure to circulating FFAs have been shown to induce insulin resistance, impair β-cell function, and compromise their survival – all pivotal processes in T2D development^5–7^. Palmitate, a main saturated FFA, has been implicated in lipotoxicity-induced T2D progression^8^. Prolonged exposure of β-cells to palmitate has been shown to impair insulin gene transcription, inhibit glucose-stimulated insulin secretion (GSIS), and induce apoptosis^6,7^. While the precise molecular mechanisms are not completely understood, mounting evidence suggests that excess FFAs or an unbalanced FFA composition can induce inflammation, endoplasmic reticulum stress, oxidative stress (with a noted increase in reactive oxygen species production), impaired autophagy, and mitochondrial dysfunction^6^. Collectively, activation of these pathways through excess FFAs such as palmitate can lead to impairments in normal β-cell functionality, insulin gene expression, apoptosis, and possibly β-cell de-differentiation^6^.

Extracellular vesicles (EV) are heterogeneous lipid membrane derived nanoparticles released by all cellular organisms. Once viewed as a means to dispose of cellular waste, research over the past three decades has recognized their role as important mediators of intercellular communication through the exchange of bioactive cargoes, including, RNAs, DNAs, proteins, and lipids^9^. Two general classes of EVs exist which are distinguished by their mode of biogenesis. Exosomes are formed through inward reverse budding of the endosomal membrane via the endocytic pathway, while ectosomes form through budding and blebbing of the plasma membrane^10^. Due to the lack of definitive biomarkers for specific EV types, broader terms are used based on biophysical and biochemical characteristics. According to the latest guidelines, there are three categories of EVs based on size exclusively – these include small EVs (50 – 150 nm in diameter), medium EVs (200 – 800 nm in diameter), and large EVs (≥ 1,000 nm in diameter)^9^.

Several lines of evidence have implicated the pancreas as a pathological tissue releasing altered EVs capable of communicating within endocrine and exocrine tissue, distal tissues/organs, and the immune system^11,12^. In both T2D and Type 1 diabetes (T1D), β-cell-derived EVs have been implicated in intra-islet communication (mainly β-cell to β-cell)^13–16^, immune cell crosstalk^17–19^, and mesenchymal stem cells^20,21^ impacting diabetes pathogenesis. Diabetogenic stress factors known to impact β-cell function, including pro-inflammatory cytokines, have also been shown to alter β-cell EV content and function through activation of specific molecular mechanisms tied to enhanced β-cell dysfunction^13,14,16^. In the context of excess FFA in T2D pathogenesis, healthy β-cell-derived EVs were shown to diminish palmitate-induced β-cell apoptosis and increase cell viability^22,23^. However, the role that lipotoxicity plays in modulating β-cell EV release, function, and mechanistic implications for T2D pathogenesis remains elusive. Therefore, in this work we demonstrate that sEVs isolated upon palmitate exposure (PAL EV) were more abundant (vs. control EVs) and induced β-cell functional failure in healthy mouse and human islets. Moreover, PAL EVs induced alterations in the islet transcriptome and showed significant lipidomic and proteomic content changes (vs. control EVs). Lastly, we implicate PAL EV-mediated activation of the TGFβ/Smad3 signaling pathway and observed that blockade of TGFβR mitigated the effects of PAL EVs on β-cell dysfunction.

## Research Design and Methods

### Animals and Study Design

12-week-old male C57BL/6L mice purchased from Jackson Labs were used for all *ex vivo* islet and high fat diet (HFD) studies. These mice were housed at the Mayo Clinic Animal Facility under standard 12 h light, 12 h dark (LD) cycle and fed either normal chow (5053; LabDiet, Inc) or high fat (60% fat, 20% protein, and 20% carbohydrates with 7% sucrose content, D12492; Research Diets, Inc). All experimental procedures and studies were approved by the Mayo Clinic Institutional Animal Care and Use Committee (IACUC).

### Cell Culture and Materials

MIN6 β-cell line was obtained from AddexBio and previously authenticated in our laboratory^13^. MIN6 cells were cultured in DMEM (high glucose) media containing 15% fetal bovine serum (FBS), 1 mM sodium pyruvate, and 50 μM β-mercaptoethanol as previously described^13^. Sodium palmitate (P9767; Sigma) was made as previously described^24^ and diluted to a final working concentration of 0.5 mM in 10% BSA diluted in 1X PBS. To generate PAL EV, MIN6 cells were treated with 0.5 mM palmitate solution for 24 h. To generate CTL EV, MIN6 cells were cultured in 10% BSA at the same calculated volume as palmitate for 24 h.

### Small EV Isolation and Characterization

#### EV isolation

sEV isolation was conducted using isolated conditioned media which was spun using differential ultracentrifugation as adapted from our previous studies^13^. In short, conditioned media was subjected to sequential spins at 2,000 x *g* for 30 minutes (2X) to rid the media of large particles and debris, filtration using a 0.2 µm filter, followed by 100,000 x *g* spins for 2 h (2X). The final pellet was resuspended in 1X PBS and aliquoted to minimize freeze-thaw cycles at -80°C.

#### NTA analysis

Nanoparticle Tracking Analysis (NTA; Malvern Panalytical) was used to quantify sEV concentration. Each fraction was diluted (1:100) in 1X PBS and ran for 5 videos, at 60 seconds each. NTA parameters were kept consistent for each run in order to make comparisons between each fraction. To obtain zeta potentials, sEV samples were diluted in dH_2_0 (1:1000) and ran on the ZetaView Particle Tracking System (Particle Metrix).

#### Transmission electron microscopy

Formvar coated 200 mesh copper grids (EMS) were stained with 1% aqueous phosphotungstic acid, pH 7.0 (EMS) to obtain transmission electron microscopy (TEM) images of sEV. Images were taken using JEOL 1400+ operating at 80kV as previously described^13^.

### Immunofluorescence Staining

For immunofluorescence staining of mouse tissue and isolated mouse islets, samples were fixed in 4% paraformaldehyde and then embedded in paraffin. Sections from isolated mouse islets treated with PAL EV and PAL EV + TGFβi (vs. UT) were stained with phospho-Smad3 (ab52903; Abcam), insulin (MAB1417; R&D Systems), and glucagon (G2654; Sigma). The relative p-Smad3 intensity was analyzed by ImageJ software and normalized by insulin expression.

### Western Blotting

For blotting of EVs, we used our previously established protocol which includes sonication of the EVs first^13^. Equal volumes of protein lysates were heated to 95°C for 5 min to denature proteins and then ran on a TGX 4-20% gradient gel (BioRad). Primary antibodies for TSG101 (PA5-82236; Invitrogen), CD9 (A1703; AbClonal), CD63 (A5271; AbClonal), Calnexin (NB100-1965; NovusBio), IAPP (A24014; AbClonal) and APP (PA1-84165; Invitrogen) were incubated overnight at 4°C. Secondary antibodies were added and images were acquired using a LiCor Odyssey Fc with Image Studio software.

### Glucose Stimulated Insulin Secretion and Islet Perifusion

C57BL/6L mouse islets were isolated as previously described via a standard collagenase protocol and recovered in RPMI 1640 media with the addition of 10% FBS and penicillin/streptomycin ^13,25^. For human studies, normal, healthy cadaveric human islets were purchased from Prodo Labs. For static GSIS, 10 islets/well (mouse) or 15 islets/well (human) were treated with 0.5 mM palmitate and ± GW4869 (5 μM; S7609, Selleckchem) for 24 h. For both mouse and human PAL EV studies, healthy islets were treated with 2X10^9^ particles each day for 48 h. For TGFβ receptor inhibition studies, LY2109761 (1 µM; S2704, Selleckchem), a TGFβ receptor I/II inhibitor was pre-treated for 1 h to mouse islets then PAL EV were added (2X10^9^ particles once a day for 48 h) or PAL EV alone were added (vs. UT islets). For all mouse and human studies, islets were subjected to a 4 mM glucose solution made in Krebs Ringer buffer (KRB) for 30 minutes, then transferred to 16 mM glucose solution made in KRB for 30 minutes.

For islet perifusion studies, isolated mouse islets were treated with PAL EV as mentioned above for 48 h (vs. UT islets). 20 islets were used to perfuse with 4 mM basal glucose and then stimulated with 16 mM glucose with the perfusate collected at 2-minute intervals^25^. For both static GSIS and islet perifusions the collected samples were run on either an ultrasensitive mouse insulin ELISA (Alpco; 80-INSMS-E10) or human insulin ELISA kit (Alpco; 80-INSHU-E10.1) and insulin values were normalized based on total insulin content.

### Lipidomic Analysis

PAL EV, CTL EV (n=4/condition), and MIN6 lysates +/- PAL treatment (n=3/condition; Supplementary Fig. 1) were isolated as described prior from MIN6 β-cells^13^. EV and cell lysates were prepared for lipidomics (CreativeProteomics) by thawing, then combining with 1.5 mL chloroform-methanol mixture (2:1, v/v). Sample mixtures were vortexed for 1 min and centrifuged for 10 min at 3,000 rpm, room temperature. The lower of the phase-separated mixture was isolated into a new tube and dried under nitrogen. Dried lipid extracts were resuspended with 200uL isopropyl alcohol-methanol mixture (1:1, v/v) and 5uL LPC(12:0) was added as internal standard. Samples were centrifuged 10 min at 12,000 rpm and 4°C, and supernatant was transferred for LC-MS analysis. Lipid separation was performed by Ultimate 3000 LC combined with Q Exactive MS (Thermo) and screened with ESI-MS. CreativeProteomics’ LC system was comprised of ACQUITY UPLC BEH C18 (100×2.1mm×1.7 μm) with Ultimate 3000 LC. The mass spectrometer was operated in positive ion mode and negative ion mode respectively with full scan MS at 70,000 resolution and data dependent MS/MS collected from 200-1200 m/z at 17,500 resolution. The electrospray ionization source was maintained at a spray voltage of 3kV at positive ion mode and -2.8kV at negative ion mode with sheath gas at 35 and auxiliary gas at 15 (arbitrary units). The inlet of the mass spectrometer was held at 350°C, and the S lens was set to 50%.

The raw data were acquired and aligned using the Lipid Search software (Thermo) based on the m/z value and the retention time of the ion signals. Ions from both ESI- or ESI+ were merged and imported into the SIMCA-P program (v.14.1) for multivariate analysis. A Principal Components Analysis (PCA) was first used as an unsupervised method for data visualization and outlier identification. Lipids were sorted by group (phospholipids, sphingolipids, etc) and by class (ceramides, cardiolipins) for comparative profiling in EVs and cell lysates, as per LipidMaps’ categorization tiers. Relative molar percentages of measured groups and classes were compared across groups (control and palmitate treated MIN6 cell lysate, EVs isolated from MIN6 cells treated with BSA and palmitate) and those with significant differences (*p* < 0.05) between EV conditions were pinpointed for further investigation for relevance in EV biogenesis and response to lipotoxic stimulus.

### RNA Isolation, Quantitative Real-time PCR, and RNA-Seq

Total RNA was extracted from C57BL/6L mouse islets using the RNeasy Plus Minikit (Qiagen, Hilden, Germany). For quantitative real-time PCR (qRT-PCR) 300 ng of mRNA was converted to complementary DNA (cDNA) using the iScript cDNA Synthesis kit (Bio-Rad, Hercules, CA). SYBR green master mix, with gene specific primers (see Supplementary Table 1) were added and qRT-PCR was performed using the Roche LightCycler 96 (Roche Diagnostics). Data was analyzed using the 2^-ΔΔCT^ method with the data normalized to housekeeping gene β-actin. For RNA-Seq analysis, total RNA was extracted from mouse islets treated with PAL EV (vs. UT) as mentioned above. RNA preparation, quality assessment, and RNA library preparation were conducted as previously noted^13^. We used *p* < 0.05 with a fold change of greater than 1.5 as a threshold for differential gene expression analysis. These genes were submitted for ontological analysis using the Database for Annotation, Visualization and Integrated Discovery (DAVID)^26^ against Kyoto Encyclopedia of Genes and Genomes (KEGG) pathways. Significantly enriched KEGG pathways (*p*<0.05) were visualized using GraphPad Prism v.9. STRING Protein-Protein Interaction Network version 11 in Cytoscape v.3.9.1 was used to assess protein-protein interactions of significantly enriched transcripts^27,28^. Gene-set enrichment analysis (GSEA) was conducted as previously noted^13,29^.

### Proteomic content of β-cell-derived lipotoxic EVs

Both PAL EV and CTL (BSA) EVs (n=3/condition) were isolated for proteomic analysis using our previously established ultracentrifugation protocol^13^. Briefly, upon addition of palmitate for 24 hours (described above), MIN6 β-cell media was collected and pre-cleared of larger particles and debris by 2x spins at 2,000 x *g* for 30 minutes each, then filtered using a 0.22 μm filter (SCGP00525; Millipore). Pre-cleared media was spun at 100,000 x *g*, for 2 hours 15 minutes, at 4°C, to pellet EVs. EVs were redispersed in PBS and spun a second time using the same settings. Supernatants were removed and samples were immediately submitted to the Mayo Clinic Proteomics Core for processing and mass spectrometry. MaxQuant v.2.3.1.0 was used for processing the raw output files, and Perseus v.1.6.5.0 for data transformation and filtering. LFQ values were used for calculations of differential protein expression between groups. Functional analysis included DAVID^26^ for association of proteins with KEGG pathways and GO terms, as well as their related p-values for significance. Enriched proteins were also submitted to PANTHER Gene Ontology for association with functional properties and visualization of term breakdowns. Z-scoring was used to visualize replicate agreement for the most significant proteins enriched in PAL- and CTL EVs, respectively. Data was graphed in GraphPad Prism v.9.

### Statistical Analysis

Statistical analyses were performed using GraphPad Prism 10 with all data presented as a mean ± SEM and where significance was denoted as *p*<0.05. For comparison of two groups, a standard Student’s t-test with Mann-Whitney post-hoc test was used and for multiple comparisons, a one-way ANOVA with Kruskal-Wallis post-hoc test.

### Data and Resource Availability

Datasets are available from the corresponding author upon reasonable request.

## Results

### Characterization of lipotoxic-induced β-cell small EVs

To generate palmitate (PAL)-exposed β-cell-derived sEV (PAL EV), MIN6 cells were treated with 0.5 mM PAL for 24 h, and sEV were isolated using our established differential ultracentrifugation procedure^13,30^. The efficacy of our sEV isolation procedure is represented in Figure 1 where Nanoparticle Tracking Analysis (NTA) was conducted on isolated untreated (control, CTL EV) MIN6 EVs and PAL EVs. A representative NTA graph depicts a significant enhancement of PAL EV (red line) compared to CTL EV (blue line) as indicated by an overall increase in concentration (particles/ml; Fig. 1A), on average an overall increase in particle concentration (∼2.4 fold; Fig. 1B), and decrease in average particle mode (Fig. 1C). Zeta potential was measured using ZetaView Multiparameter NTA (Particle Metrix) to assess the surface charge of the EVs which typically harbor a net negative charge under normal physiological states^31^. Interestingly, it was noted that PAL EVs were more negatively charged compared to CTL EVs (Fig. 1D), suggesting that pathological PAL EVs possess enhanced stability and potential efficacy in intercellular communication. Both PAL EV and CTL EV (with MIN6 cell lysate as a control) were subjected to western blot analysis of known sEV biogenesis markers TSG101, CD9, and CD63, with the absence of Calnexin (negative control for EVs) in both EV samples, validating EV purity (Fig. 1E). Transmission electron microscopy (TEM) of PAL EV displayed intact vesicles within the typical size range of sEV (scale bar represents 100 nm; Fig. 1F). Lipidomic analysis was conducted to determine EV lipid alterations between PAL EVs and CTL EVs (n=4/group) vs. total MIN6 cell lysate +/- palmitate treatment (n=3/group). A comparison of the total number of unique lipid species identified in MIN6 cell lysates (882) and CTL EVs (276) with 161 lipids that overlap was done (Fig. 1G). In addition, major lipid groups as a molar percentage of total lipids were identified in MIN6 control cell lysate (+BSA) and MIN6 cells treated with 0.5 mM palmitate (24h; Supplementary Fig. 1). Although there were no indicated changes in the major lipid groups, alterations in differentially expressed lipid species in each group were noted in the bar graphs after palmitate addition ((FC)>1; Supplementary Fig. 1). However, more significant alterations in lipid composition were noted in PAL– vs. CTL EVs including enrichment of phosphatidylethanol, monoglyceride, phosphatidylserine, and hexosylceramides (Fig. 1H).

**Figure 1:**
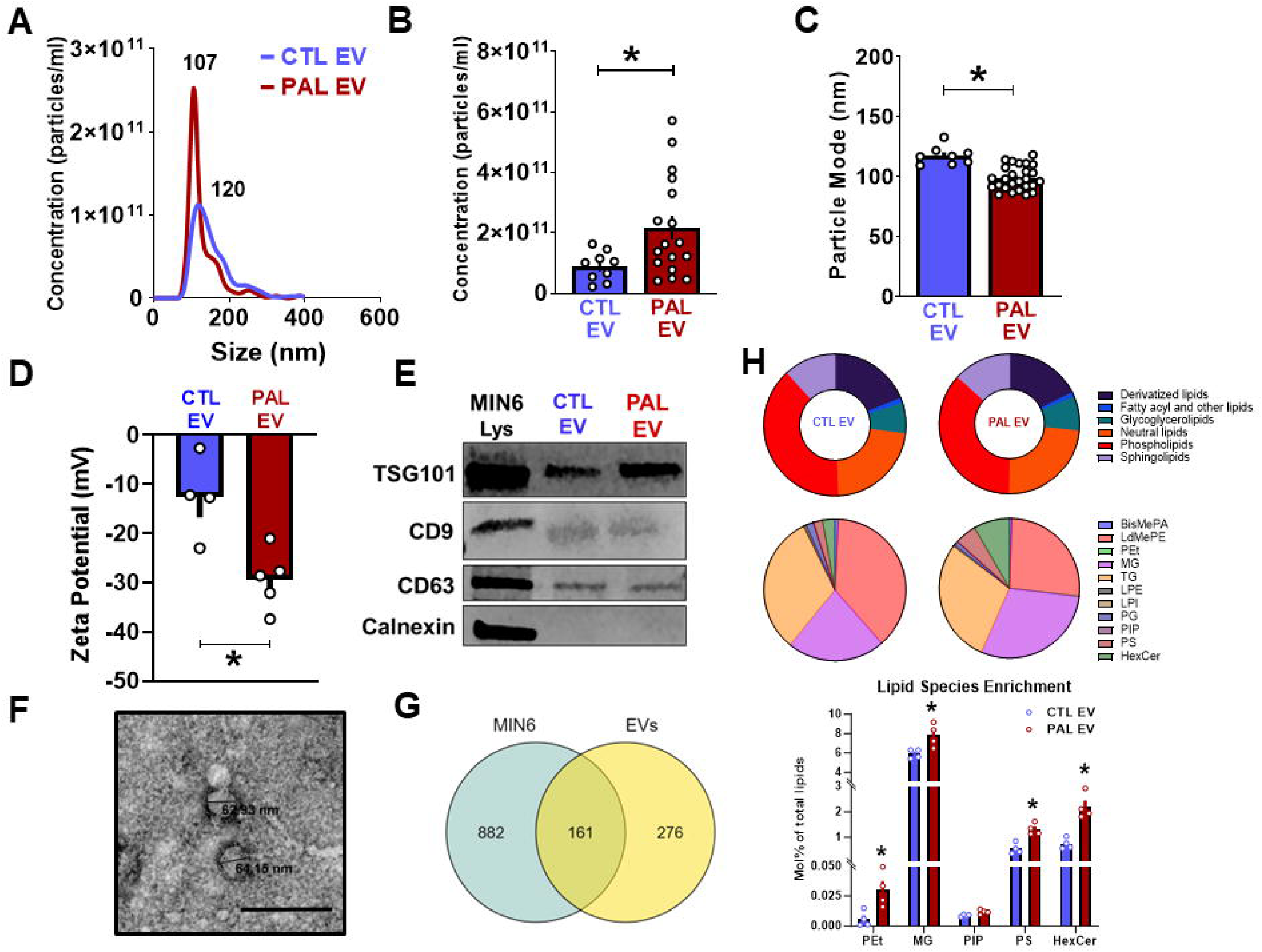
Characterization of lipotoxicity-induced MIN6 mouse β-cell small EVs. *A:* Mouse MIN6 β-cell line was treated with palmitate (PAL) or control (BSA) for 24 h and sEV were isolated from the conditioned media. Overlap of a representative Nanoparticle Tracking Analysis (NTA) graph depicting untreated (BSA) control (CTL)EV in blue and palmitate (PAL)EV in red with the mode for each (120 nm and 110 nm, respectively). *B and C*: Overall quantification of particles released from CTL-(n=9 independent isolations) and PAL EV (n=15 independent isolations) and average mode of those particles (*C*). *D*: Zeta potential was acquired from CTL EV (n=4) and PAL EV (n=5) using ZetaView (Particle Matrix). *E*: Western blot analysis of CTL EV, PAL EV, and MIN6 lysate (control) for sEV biogenesis markers TSG101, CD9, and CD63, with Calnexin as a negative control (representative example from n=4-10 EV blots. *F*: Transmission electron microscopy (TEM) of isolated PAL EV (representative example from n=3 EV isolations); scale bar represents 100 nm. *G*: Venn diagram depicting unique and overlapping lipid specifies from lipidomic analysis of MIN6 EVs (n=4/condition) vs. MIN6 lysate (n=3/condition). *H*: Differential expression of lipid species defined as a molecular percentage of total lipids from PAL EV vs. CTL EV. Values are a mean ± SEM. Statistical significance among groups is indicated by *, *p*<0.05.

A comprehensive analysis of the proteomes of both CTL- and PAL EVs was conducted. The volcano plot revealed 91 differentially expressed proteins in PAL EV vs. 67 in CTL EVs (Fig. 2A) and of those proteins 70 were only present in either PAL- or CTL EVs with 1,397 proteins in common (Log_2_ (FC)>1; Fig. 2B). Enriched proteins in PAL EV include Nuclear casein kinase and cyclin-dependent kinase substrate 1 (NUCKS1), a nuclear protein involved in chromatin remodeling and regulation of gene transcription; nardilysin (NRD1), a N-arginine dibasic convertase; and amyloid precursor protein (APP), the precursor molecule to amyloid beta (Aβ) (Fig. 2C). Moreover, using PANTHER GO Slim analysis for Protein classes several proteins were enriched in PAL EV that are associated with β-cell regulation and function including islet amyloid polypeptide (IAPP), insulin 2 (INS2), and previously mentioned APP (Fig. 2D). Expression of these proteins were confirmed through western blotting of PAL EVs (vs. CTL EVs and MIN6 lysate) for IAPP (Fig. 2E), and APP (Fig. 2F).

**Figure 2:**
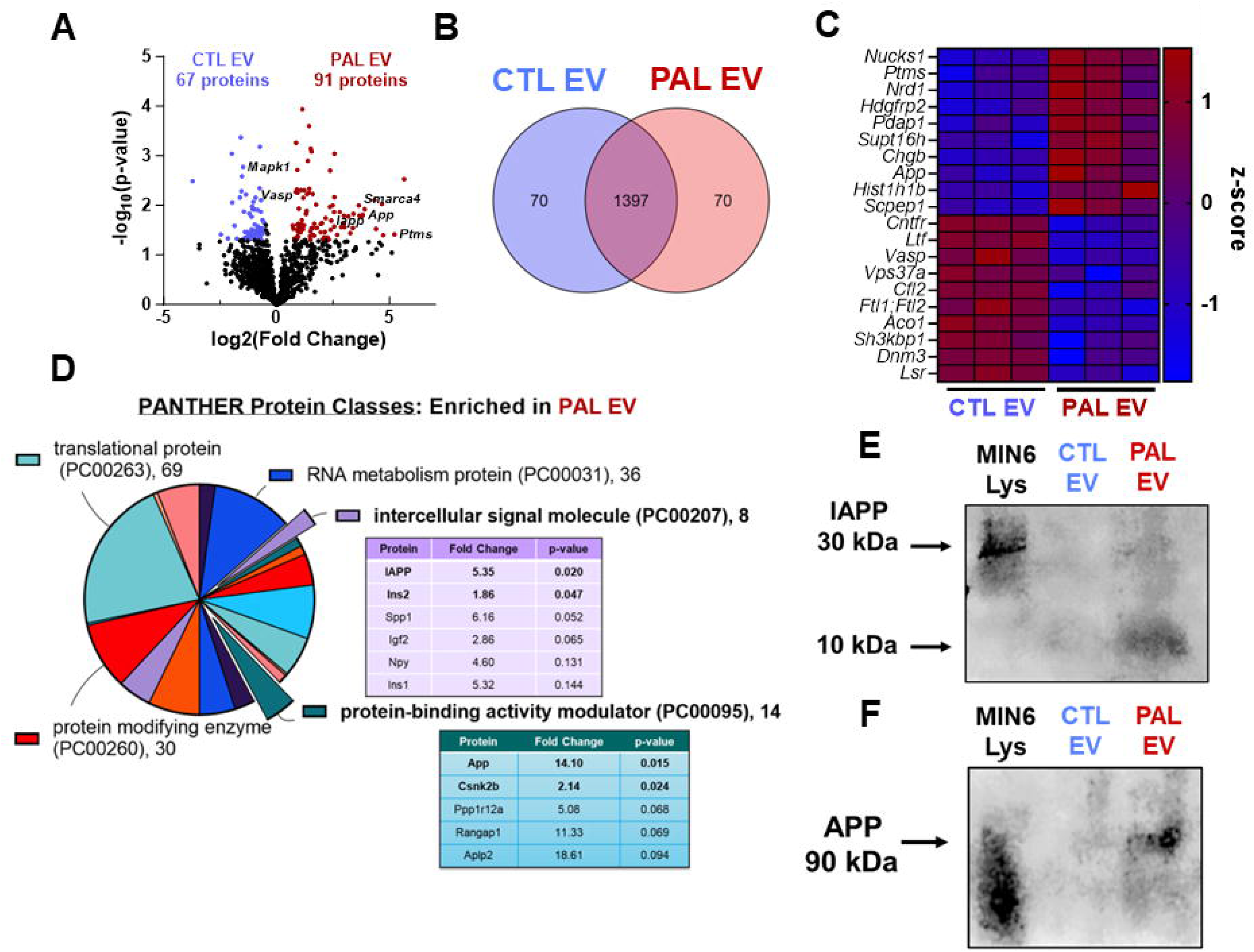
Proteomic analysis of lipotoxic β-cell small EV. *A*: PAL EV and CTL EV were isolated upon PAL or BSA (control) treatment on MIN6 β-cell line for 24 h. Final sEV pellets were subjected to proteomic analysis. Volcano plot depicting differentially expressed proteins in PAL EV vs. CTL EV (n=3/condition; FC>1.5; *p*<0.05). *B*: Venn diagram revealed 70 uniquely enriched proteins in both CTL- and PAL EV with ∼1400 overlapping proteins. *C*: Heatmap shows top 10 upregulated and 10 downregulated proteins found in PAL EVs vs. CTL EV. *D*: Panther Go Slim analysis of protein classes that were enriched in PAL EV are depicted in the pie chart. Inserts reveal differentially expressed proteins for “intercellular signal molecule” and “protein-binding activity modulator along with FC and p value. *E-F*: Western blot confirmation of differentially expressed proteins, IAPP and APP relating to β-cell function and identity in PAL EV (vs. CTL EV an MIN6 lysate).

### Small EVs contribute to lipotoxic-mediated β-cell dysfunction

To determine if lipotoxicity-induced sEV release impacts β-cell function, we used GW4869, an inhibitor of sEV biogenesis, which inhibits the formation of ceramide-mediated intraluminal vesicle formation in multivesicular bodies^13,32^. MIN6 cells were treated with PAL ± GW4869 (5 μM) for 24 h and sEV release was quantified using NTA analysis. sEV biogenesis blockade significantly reduced PAL EV release by ∼84% with GW4869 addition (Fig. 3A) and reduced the overall particle mode (Fig. 3B). As previously noted, palmitate (PAL) addition to isolated C57BL/6L mouse islets induced β-cell dysfunction as noted by significant impairments in glucose stimulated insulin secretion (GSIS; Fig. 3C) with a ∼63% decrease in insulin stimulation index (SI) compared to untreated islets (UT; Fig. 3D). However, inhibition of sEV biogenesis with PAL addition, strikingly improved GSIS (∼5 fold), as depicted in the SI graph with no alterations in GSIS with GW4869 alone, suggesting that lipotoxic sEVs are potential contributors to β-cell dysfunction under lipotoxicity (Fig. 3C-D). Moreover, to determine if blockade of sEV biogenesis also improves GSIS in islets isolated from mice exposed to high fat diet (HFD), C57BL/6L male mice were put on a HFD or chow diet for 10 weeks and isolated islets were exposed to GW4869. Consistently, administration of GW4869 enhanced insulin stimulation index (p<0.05, Supplementary Fig. 2). Similarly, to assess if lipotoxicity-induced sEV also impact β-cell function in human islets, we treated healthy human cadaveric islets with PAL or PAL+GW4869 compound for 24 hours in the same manner as we did for the mouse islet experiments. Results showed PAL addition induced β-cell dysfunction as previously observed (∼82% decrease in SI; Fig. 3E), however upon sEV biogenesis inhibition, we noted a significant restoration of GSIS (∼5.8 fold increase in SI vs. PAL only; *p* < 0.05; Fig. 3F).

**Figure 3.**
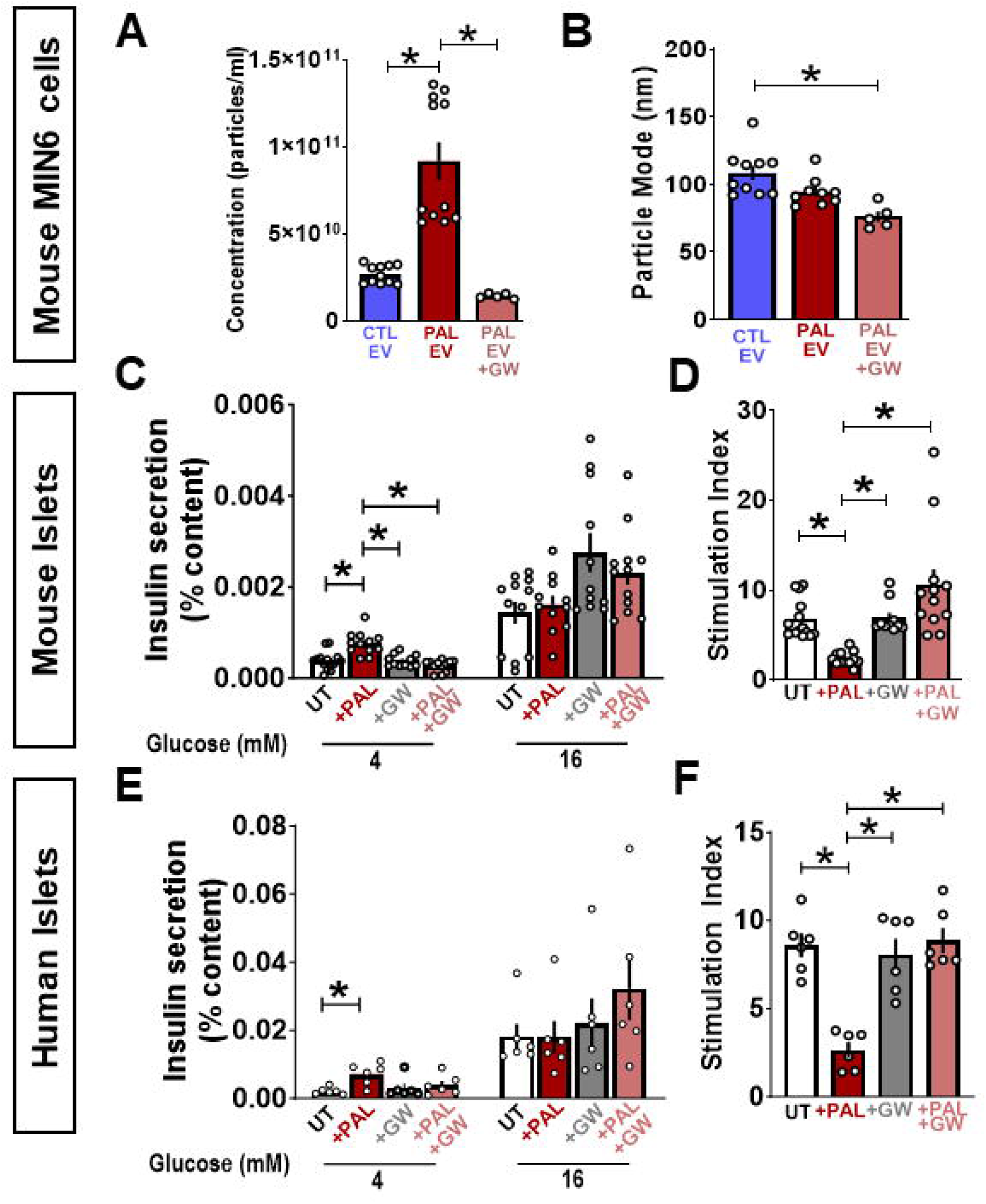
Small EV generation contributes to lipotoxic-mediated β-cell dysfunction. *A and B*: MIN6 cells were treated with PAL or PAL+GW4869 (5 μM; 24h) vs. Control (BSA) EV and EV particle concentrations (A) and mode (B) were assessed using NTA (n=5-11 independent EV isolations). *C and D*. C57BL/6L mouse islets were treated with palmitate (PAL; 0.5 mM) ± GW4869 for 24 h. Static glucose stimulated insulin secretion (GSIS) was assessed at 4 mM basal and 16 mM stimulatory glucose concentrations and insulin stimulation index is expressed as 16 mM glucose divided by 4 mM basal concentrations (n=10-14 independent experiments per condition). *E and F*. Healthy human islets were treated with 0.5 mM PAL ± GW4869 (5 μM) for 24 h and static GSIS was conducted. Insulin stimulation index is depicted as 16 mM stimulatory values divided by 4 mM basal values (n=6 independent experiments per condition).

### Lipotoxic β-cell small EVs contribute to β-cell dysfunction in isolated islets

To determine if PAL EV have the propensity to impact β-cell function, C57BL/6L mouse islets were isolated and exposed to PAL EV or CTL EV (vs. untreated islets) for 48 hours. Islets were subjected to *in vitro* static GSIS which resulted in a significant increase in 4 mM basal glucose values and reduction in 16 mM glucose with PAL EV treatment in comparison to CTL EV addition and untreated islets (*p*<0.05; Fig. 4A). Additionally, insulin stimulation index revealed an ∼80% decrease in GSIS response with PAL EV addition compared to untreated islets (Fig. 4B). To corroborate the static GSIS experiment, an *in vitro* dynamic perifusion assay was conducted on C57BL/6L mouse islets exposed to PAL EV (vs. untreated islets). The results showed that PAL EV reduced 16 mM stimulatory values, suggesting the capacity of PAL EV to attenuate β-cell function (Fig. 4C). Moreover, C57BL/6L mouse islets exposed to PAL EV significantly diminished expression of key β-cell identity markers, *Ins1*, *Ins2*, and *Pdx1* in (Supplementary Fig. 3A; *p* < 0.05). In addition, we noted that antecedent exposure of PAL EV to C57BL/6L mouse islets did not impact β-cell apoptosis or proliferation (Supplementary Fig. 3B-C). Lastly, we exposed healthy human cadaveric islets to PAL (0.5 mM; 24 h) and sEVs were isolated using differential ultracentrifugation to generate human PAL EV (hPAL EV). Particles were quantified using NTA shown in the representative graph with the average particle concentration and zeta potential (Fig. 4D), and expression of sEV biogenesis markers (with absence of Calnexin; Fig. 4E). hPAL EVs were used to treat healthy human cadaveric islets for 48 h then subjected to static GSIS which revealed apparent β-cell dysfunction as noted by an overall reduction in insulin SI (∼38%; Fig. 4F-G).

**Figure 4:**
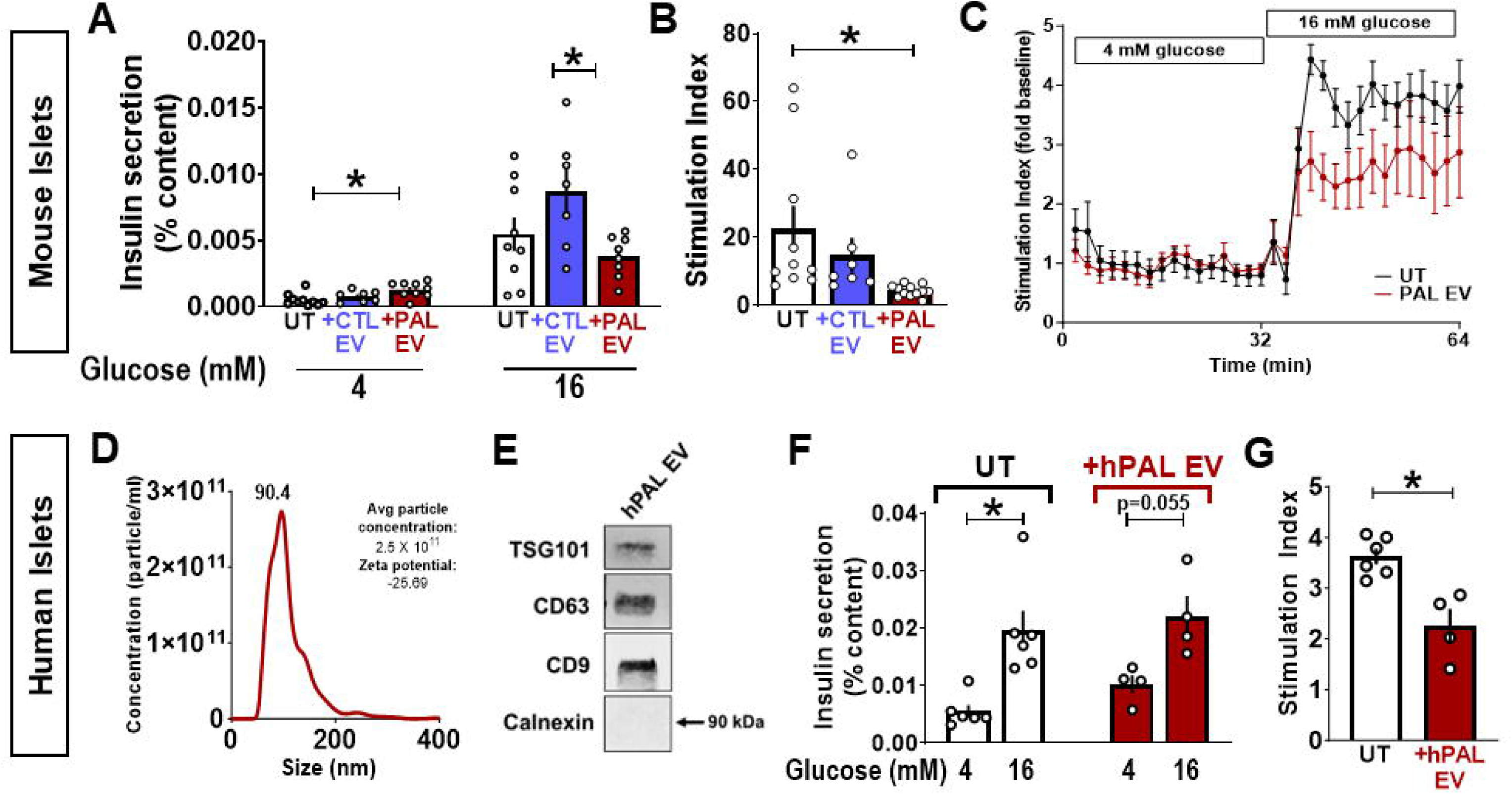
Lipotoxic-induced β-cell small EVs induce β-cell dysfunction. *A and B*: C57BL/6L mouse islets were treated with 2X10^9^ particles (either CTL- or PAL EV) each day for 48 h. Static GSIS was conducted along with determination of insulin stimulation index (n=7-12 independent experiments per condition). *C*: For islet perifusion, C57BL/6L mouse islets were treated with 2X10^9^ particles each day for 48 h (vs. UT islets) and islet perifusion was performed at 4 mM basal glucose (0-32 min) and 16 mM glucose (32-62 min). Samples were taken at 2 min intervals with a n=3-5 independent experiments per condition. *D*: Representative NTA graph for hPAL EV depicting concentration (particles/ml) by size (nm), average particle concentration, and zeta potential (n=2 isolations conducted). *E*. Western blot analysis depicting expression of sEV biogenesis markers TSG101, CD9, CD63 in a representative hPAL EV sample with the absence of Calnexin. *F and G*: Static GSIS of hPAL EV exposure to healthy human islets with n=4-6 independent experiments per condition. Values are a mean ± SEM. Statistical significance among groups is indicated by *, *p*<0.05.

### Lipotoxic β-cell-derived small EVs alter the islet transcriptional landscape

To understand the mechanistic alterations that occur with PAL EV addition to isolated mouse islets (vs. untreated islets), we conducted global RNA-Sequencing analysis. Our analysis revealed ∼880 genes uniquely enriched with PAL EV addition to islets, while over 1,000 genes were unique to control (untreated) islets (*p* < 0.05, with fold change (FC) ≥ 1.5; Fig. 5A). The top 10 most upregulated genes with PAL EV addition vs. untreated islets include *Snurf* (9.6 fold), *Tff2* (8.2 fold), and *Reg3g* (8.2 fold) (*p* < 0.05; Fig. 5B). Additionally, KEGG pathway analysis highlighted the enrichment of genes that are associated with pathways such as Protein digestion and absorption, ECM receptor interaction, and Pancreatic secretion. Conversely, downregulated pathways included Protein processing in endoplasmic reticulum, HIF-1 signaling, and Protein export (*p* < 0.05; Fig. 5C). The interactome depicts pathway interactions from enriched gene transcripts (*p* < 0.05; Fig. 5D) and additional Gene Ontology (GO) term analysis for Cellular Component (Fig. 5E), Biological Processes (Fig. 5F), and Molecular Function (Supplementary Fig. 4) showed enrichment of genes associated with extracellular matrix and reorganization pathways, while downregulated pathways included processes with the endoplasmic reticulum (ER).

**Figure 5:**
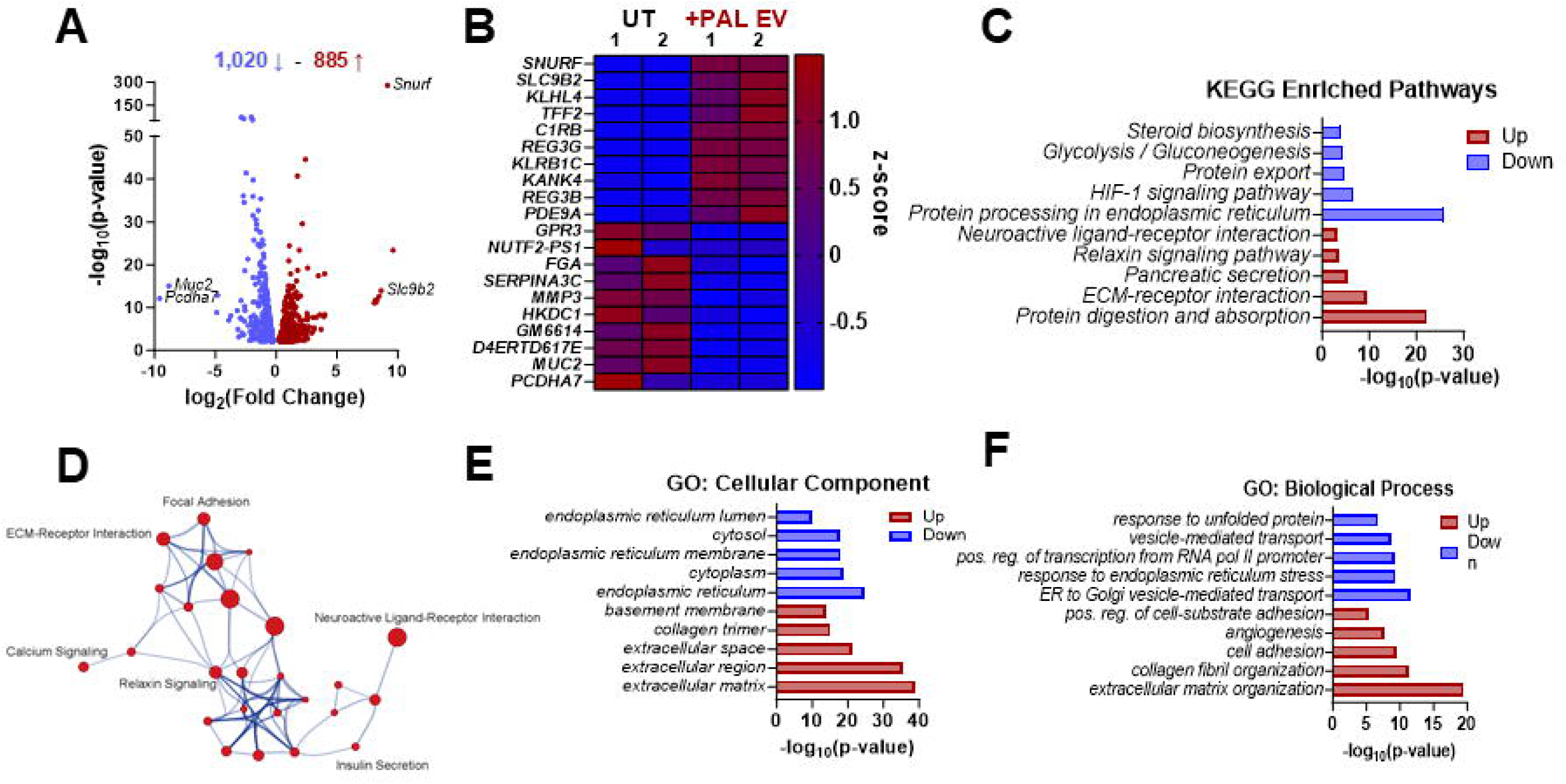
Islet transcriptome alterations upon antecedent exposure to lipotoxic β-cell small EVs. *A*: C57BL/6L mouse islets were exposed to PAL EV (2X10^9^ EV/day; 48 h) vs. UT islets (n=2 biological repeats per condition). RNA-sequencing was conducted, and volcano plot depicts differentially expressed genes (FC>1.5; *p*<0.05) where 885 genes were found to be upregulated and ∼1,000 genes downregulated upon PAL EV addition to islets. *B*. Heatmap shows normalized topmost upregulated and downregulated transcripts in PAL EV treated vs. UT islets. *C*: KEGG pathway analysis was conducted showing top 4 most upregulated and downregulated pathways (*p* < 0.05). *D*: Significantly enriched KEGG pathways (*C*) show interactions through *in silico* pathway analysis such as Focal Adhesion and ECM Receptor Interaction. Size of the node denotes the number of transcripts associated with the pathway. *E-F*: Gene ontology (GO) analysis for Cellular Component and Biological Processes reveal both up- and downregulated gene transcripts associated with ECM organization (up) and endoplasmic reticulum processes (down) (*p* < 0.05).

### Lipotoxic β-cell small EV activate the TGFβ/Smad3 pathway during β-cell dysfunction

To determine the molecular insights of PAL EV-mediated transcriptional alterations in islets, we employed *de novo* motif analysis revealing differentially enriched transcription factor binding sites in promoter regions of genes enriched in islets exposed to PAL EV (Fig. 6A). Interestingly, the top significantly enriched transcription factor binding sites mapped to *Smad* genes, including *Smad2*, *-3*, and -*4* which are involved in the TGFβ/Smad3 pathway (Fig. 6A). We used gene set enrichment analysis (GSEA) to determine if there was any significant enrichment of genes relating to the TGFβ/Smad3 pathway. Indeed, we found enrichment of genes associated with TGFβ receptor binding pathway upon PAL EV addition to islets (Fig. 6B). The heatmap depicts all the genes enriched in the TGFβ receptor binding pathway which include several of the TGFβ ligands (*e.g. Tgf*β*1*, *Tgf*β*2*, *Tgf*β*3*), TGFβ receptors (*e.g. Tgf*β*R2*, *Tgf*β*R3*), and Smad gene (*Smad3*) (Fig. 6C).

**Figure 6:**
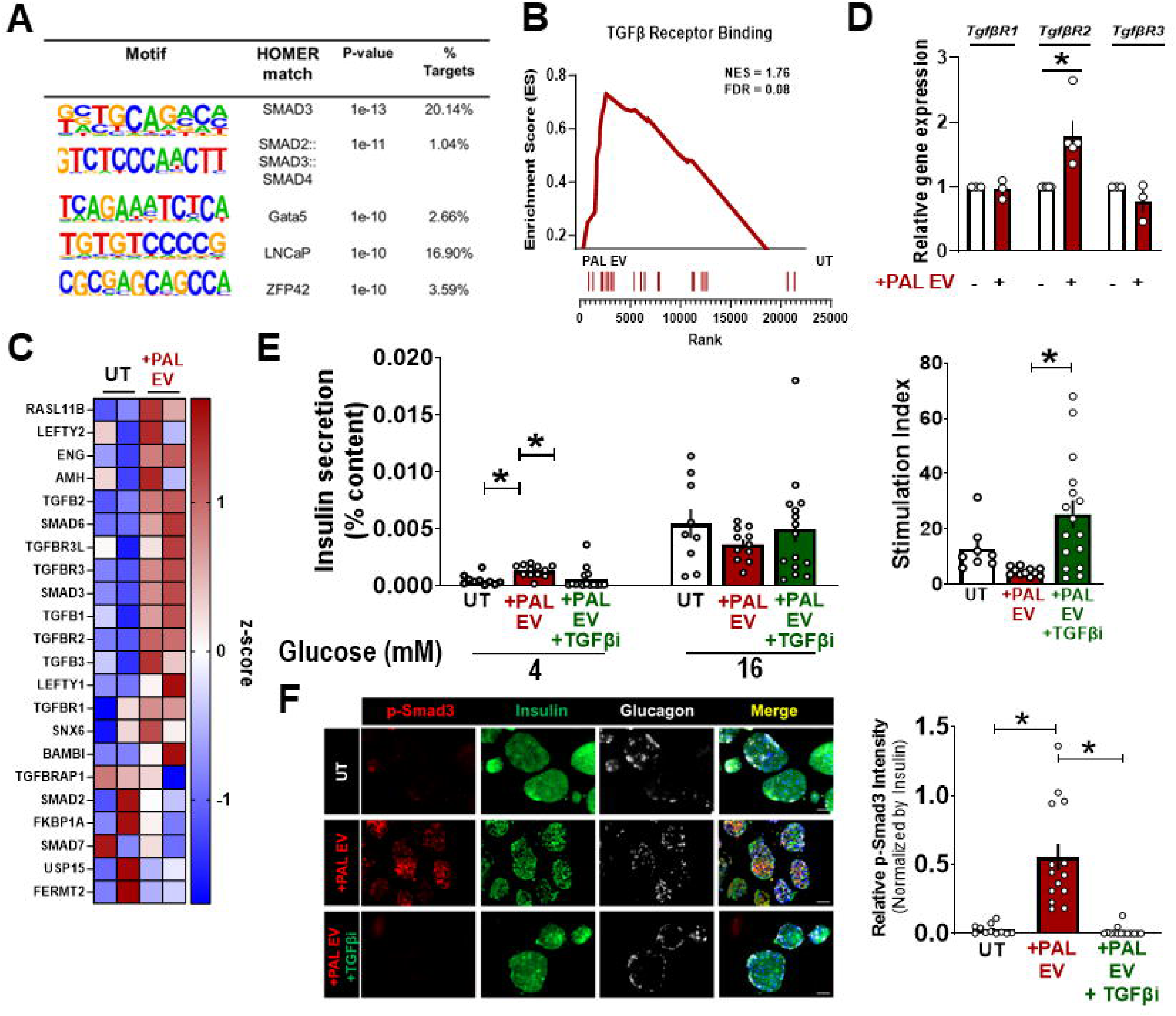
Lipotoxic β-cell-derived small EV activate the TGFβ-Smad3 pathway. *A*: *De novo* motif analysis using HOMER was used to assess transcription factor motifs that were enriched in upregulated genes found in the RNA-Seq data (PAL EV vs. UT). Topmost enriched transcription factor binding sites in PAL EV treated mouse islets were those from the TGFβ/Smad3 pathway including: Smad2, Smad3, Smad4 (*p* < 0.05). *B*: Gene Set Enrichment Analysis (GSEA) was conducted on a subset of genes that regulate TGFβ Receptor Binding (FDR=0.08) that was found to be enriched in PAL EV treated islets (vs. UT). *C*: Heatmap depicts the specific enriched gene transcripts in PAL EV treated islets that are associated with TGFβ Receptor Binding from the GSEA (*B*). *D*: Gene expression analysis using qRT-PCR to confirm expression of *Tgf*β*r1*, *Tgf*β*r2*, and *Tgf*β*r3* (n=3-5 independent experiments per condition). *E*: C57BL/6L mouse islets were exposed to PAL EV (2X10^9^ EV/day; 48 h) or PAL EV + TGFβi (TGFβ receptor I/II inhibitor (1 μM; LY2109761)); vs. UT islets. Static GSIS was conducted along with insulin stimulation index (n=8-16 independent experiments conducted per condition). *F*: Immunofluorescence staining was conducted on mouse islets exposed to PAL EV (2X10^9^ EV/day; 48 h) or PAL EV + TGFβi (TGFβ receptor I/II inhibitor (1 µM; LY2109761)); vs. UT islets. Phospho-Smad3 expression was noted in the red, insulin in green, and glucagon in white. Relative p-Smad3 intensity was quantified using ImageJ software and normalized to insulin expression. Values are a mean ± SEM. Statistical significance among groups is indicated by *, *p*<0.05.

Enhanced expression of TGFβR2 gene expression with PAL EV addition was confirmed via qRT-PCR (Fig. 6D). Therefore, to confirm if PAL EV are responsible for activation of the TGFβ/Smad3 pathway, we used a pharmacological inhibitor specific to TGFβR1/2 (1 µM; LY2109761, TGFβi) to block pathway activation. C57BL/6L mouse islets were exposed to PAL EVs or PAL EVs + TGFβi and GSIS was conducted (vs. UT islets; Fig. 6E). TGFβ receptor blockade in the presence of PAL EVs significantly improved GSIS, as evidenced by a ∼5.5 fold increase in SI (vs. PAL EV only; Fig. 6E). Activation of the TGFβ/Smad3 pathway was further confirmed through immunofluorescence staining of C57BL/6L mouse islets treated with PAL EV or PAL EV+ TGFβi (vs. UT islets). pSmad-3 (red) expression was used to delineate TGFβ/Smad3 pathway activation. Quantification of relative p-Smad3 intensity showed a significantly increase upon PAL EV addition and decreased upon PAL EV+ TGFβi treatment (vs. PAL EV alone; Fig. 6F).

## Discussion

Regulation of glucose homeostasis relies on inter-organ communication between the pancreas and other metabolically active tissues. Diabetogenic stressors (*i.e.* inflammation, excess FFA, glucotoxicity, etc) have been shown to enhance dysregulation of this communication through the release of signaling factors to impede healthy β-cell function and T2D progression. Growing evidence has implicated EVs as novel signaling effectors in the pathogenesis of diabetes through their ability to induce molecular alterations in β-cells, immune cells, and other metabolically active tissues. Here, we completed a full characterization of lipotoxic-induced β-cell sEVs through assessment in alterations in concentration, lipid and protein content, and function. We used both mouse and human islet models to determine the capacity of PAL EV to induce β-cell dysfunction and enact changes in global islet gene expression. Moreover, we provide evidence of the molecular mechanistic alterations imparted by PAL EV through activation of the TGFβ/Smad3 pathway where TGFβ receptor blockade improved overall β-cell function in the presence of PAL EV.

Studies have reported that islet exposure to FFA palmitate induces ceramide generation and subsequently inhibits insulin gene expression and β-cell apoptosis^33–35^. Mechanistically, ceramides have been shown to impact these processes through direct activation of apoptotic pathways and/or induction of ER stress^36^. Although 24 h exposure to 0.5 mM palmitate did not significantly alter any particular lipid species in MIN6 cells, we did see significant alterations in PAL EVs vs. CTL EVs including enrichment in hexosylceramides. Alterations in the lipid composition has been shown to change membrane fluidity and curvature with ceramides having a critical role in membrane curvature and thus the formation of sEV (*i.e.* intraluminal vesicles) of MVBs. Studies have shown that palmitate exposure induced lipotoxic ER stress, leading to ceramide rich EV release from hepatocytes^37–39^. Thus, it is plausible that ceramide enrichment under lipotoxic stress in β-cells may also play a vital role in the enhancement of sEV production as reported in our study. Moreover, as the distribution of sEV membrane lipids has been shown to impact sEV generation, release, and targeted delivery to nearby cells/tissues, alterations in intercellular homeostasis can affect overall sEV contribution. Thus, understanding the key alterations in PAL EV vs. CTL EV lipid composition could shed light on their mechanistic role as contributors to β-cell dysfunction. For example, phosphatidylserine (PS) is found typically on the inner leaflet of the lipid bilayer but becomes “exposed” or externalized during apoptosis which signals recruitment of macrophages for destruction^40,41^. Similarly, EVs have also been shown to display PS on their outer membrane^42,43^ which has been shown to be enhanced during disease pathology such as tumorigenesis^44–47^. As our data indicates an enrichment of PS on PAL EV (vs. CTL EV), it may be plausible that these sEV expose PS on the outer leaflet thus enhancing β-cell susceptibility to immune cell attack. Further investigation will go into understanding these lipid alterations due to PAL exposure and in addition, the contribution of other pathologically relevant FFAs that have been implicated in T2D^48^.

Our RNA-Seq data of islets exposed to PAL EV revealed significant enrichment of genes associated with extracellular matrix (ECM) remodeling, however the connection between lipotoxic EV exposure, activation of the TGFβ/Smad3 pathway, and ECM remodeling requires further investigation. The ECM which consists of a large network of proteins that support the structure and communication between the cells/tissues, is also adaptive in response to cellular stressors such as inflammation and injury^49^. Pathological remodeling of the ECM has been noted in response to overnutrition in obesity where ECM expansion was shown to induce membrane stiffness and promote insulin resistance^50^. Moreover, a direct connection between ECM remodeling and TGFβ signaling exists where pro-fibrogenic responses enhance ECM deposition through TGFβ regulatory mechanisms^51^. TGFβ signaling has been shown to play a significant role in the developing pancreas^52–54^, however several studies have shed light on its role in the adult pancreas. For example, *Smad3* was shown to bind to the insulin gene promoter and suppress transcription while *Smad3* deficient mice showed only mild hyperglycemia with improvements in glucose tolerance and GSIS^55^. Moreover, small molecule inhibition of the TGFβ pathway was shown to be protective against β-cell de-differentiation and restored overall β-cell identity^56^ and promote β-cell replication^57^. Interestingly, our proteomic analysis of PAL EV showed enrichment of β-cell regulatory and functional genes IAPP, INS2, and APP. Thus, future work will be conducted on understanding the link between PAL EV-mediated ECM remodeling and activation of the TGFβ pathway via altered PAL EV content to induce β-cell dysfunction.

In conclusion, the present study offers insight into the molecular underpinnings of lipotoxic-stress mediated β-cell dysfunction through the auspices of molecular mediators, sEV which under these conditions show altered content and function. We show that palmitate-induced stress enhances sEV secretion from β-cells (PAL EV) and alters both the overall lipid and proteomic content (vs. CTL EV). sEV inhibition not only reduces overall PAL EV formation/release, but also improves lipotoxic β-cell dysfunction *ex vivo* in mouse and human islets, suggesting the importance of sEV as mediators of lipotoxic-induced β-cell stress. In addition, mouse islet exposure to PAL EV induced transcriptional alterations centering around ECM interactions and remodeling. Moreover, our RNA-Seq data also revealed a mechanistic link between PAL EV addition and activation of the TGFβ/Smad3 pathway in islets where receptor inhibition of the TGFβ I/II receptor improved PAL EV-mediated β-cell dysfunction. Future directions will involve deciphering more in-depth molecular mechanisms of PAL EV-mediated β-cell dysfunction with the long-term goal of developing novel sEV-based strategies to protect or enhance the loss of β-cell functional mass during the pathogenesis of T2D.

## Acknowledgments

The authors thank Charles Howe, PhD and Ngoc Hoan (Henry) Nguyen, PhD from Mayo Clinic for their assistance in running the ZetaView Nanoparticle Tracking Analyzer.

## Funding

This study was supported by the National Institutes of Health (NIH) grants K01 DK129208 (N.J.) and JDRF 2-SRA-2022-1272-S-B (N.J.).

## Duality of Interest

No potential conflicts of interest relevant to this article were reported.

## Author Contribution

N.J. and A.V.M. conceived and designed the study. A.R., A.H., T.K.H., G.A., K.S.R., A.S.G., performed experiments. A.R., A.H., M.R.B., analyzed data. N.J. wrote the manuscript. All authors reviewed the manuscript.

## Notes

### Competing Interest Statement

The authors have declared no competing interest.

## References

1 Kahn, S. E., Cooper, M. E. & Del Prato, S. Pathophysiology and treatment of type 2 diabetes: perspectives on the past, present, and future. Lancet 383, 1068–1083 (2014). 10.1016/S0140-6736(13)62154-6

2 Kahn, S. E., Zraika, S., Utzschneider, K. M. & Hull, R. L. The beta cell lesion in type 2 diabetes: there has to be a primary functional abnormality. Diabetologia 52, 1003–1012 (2009). 10.1007/s00125-009-1321-z

3 Galicia-Garcia, U. et al. Pathophysiology of Type 2 Diabetes Mellitus. Int J Mol Sci 21 (2020). 10.3390/ijms21176275

4 Boden, G. Obesity, insulin resistance and free fatty acids. Curr Opin Endocrinol Diabetes Obes 18, 139–143 (2011). 10.1097/MED.0b013e3283444b09

5 Ye, R., Onodera, T. & Scherer, P. E. Lipotoxicity and beta Cell Maintenance in Obesity and Type 2 Diabetes. J Endocr Soc 3, 617–631 (2019). 10.1210/js.2018-00372

6 Lytrivi, M., Castell, A. L., Poitout, V. & Cnop, M. Recent Insights Into Mechanisms of beta-Cell Lipo- and Glucolipotoxicity in Type 2 Diabetes. J Mol Biol 432, 1514–1534 (2020). 10.1016/j.jmb.2019.09.016

7 Poitout, V. et al. Glucolipotoxicity of the pancreatic beta cell. Biochim Biophys Acta 1801, 289–298 (2010). 10.1016/j.bbalip.2009.08.006

8 Oh, Y. S., Bae, G. D., Baek, D. J., Park, E. Y. & Jun, H. S. Fatty Acid-Induced Lipotoxicity in Pancreatic Beta-Cells During Development of Type 2 Diabetes. Front Endocrinol (Lausanne) 9, 384 (2018). 10.3389/fendo.2018.00384

9 Welsh, J. A. et al. Minimal information for studies of extracellular vesicles (MISEV2023): From basic to advanced approaches. J Extracell Vesicles 13, e12404 (2024). 10.1002/jev2.12404

10 Buzas, E. I. The roles of extracellular vesicles in the immune system. Nat Rev Immunol 23, 236–250 (2023). 10.1038/s41577-022-00763-8

11 Noren Hooten, N. & Evans, M. K. Extracellular vesicles as signaling mediators in type 2 diabetes mellitus. Am J Physiol Cell Physiol 318, C1189–C1199 (2020). 10.1152/ajpcell.00536.2019

12 Javeed, N. Shedding Perspective on Extracellular Vesicle Biology in Diabetes and Associated Metabolic Syndromes. Endocrinology 160, 399–408 (2019). 10.1210/en.2018-01010

13 Javeed, N. et al. Pro-inflammatory beta cell small extracellular vesicles induce beta cell failure through activation of the CXCL10/CXCR3 axis in diabetes. Cell Rep 36, 109613 (2021). 10.1016/j.celrep.2021.109613

14 Guay, C., Menoud, V., Rome, S. & Regazzi, R. Horizontal transfer of exosomal microRNAs transduce apoptotic signals between pancreatic beta-cells. Cell Commun Signal 13, 17 (2015). 10.1186/s12964-015-0097-7

15 Ribeiro, D. et al. Extracellular vesicles from human pancreatic islets suppress human islet amyloid polypeptide amyloid formation. Proc Natl Acad Sci U S A 114, 11127–11132 (2017). 10.1073/pnas.1711389114

16 Lakhter, A. J. et al. Beta cell extracellular vesicle miR-21-5p cargo is increased in response to inflammatory cytokines and serves as a biomarker of type 1 diabetes. Diabetologia 61, 1124–1134 (2018). 10.1007/s00125-018-4559-5

17 Cianciaruso, C. et al. Primary Human and Rat beta-Cells Release the Intracellular Autoantigens GAD65, IA-2, and Proinsulin in Exosomes Together With Cytokine-Induced Enhancers of Immunity. Diabetes 66, 460–473 (2017). 10.2337/db16-0671

18 Guay, C. et al. Lymphocyte-Derived Exosomal MicroRNAs Promote Pancreatic beta Cell Death and May Contribute to Type 1 Diabetes Development. Cell Metab 29, 348–361 e346 (2019). 10.1016/j.cmet.2018.09.011

19 Giri, K. R. et al. Molecular and Functional Diversity of Distinct Subpopulations of the Stressed Insulin-Secreting Cell’s Vesiculome. Front Immunol 11, 1814 (2020). 10.3389/fimmu.2020.01814

20 Chen, J. et al. Mesenchymal stem cell-derived exosomes protect beta cells against hypoxia-induced apoptosis via miR-21 by alleviating ER stress and inhibiting p38 MAPK phosphorylation. Stem Cell Res Ther 11, 97 (2020). 10.1186/s13287-020-01610-0

21 Nojehdehi, S. et al. Immunomodulatory effects of mesenchymal stem cell-derived exosomes on experimental type-1 autoimmune diabetes. J Cell Biochem 119, 9433–9443 (2018). 10.1002/jcb.27260

22 Tang, S. et al. Neutral Ceramidase Secreted Via Exosome Protects Against Palmitate-Induced Apoptosis in INS-1 Cells. Exp Clin Endocrinol Diabetes 125, 130–135 (2017). 10.1055/s-0042-116314

23 Sun, Y., Mao, Q., Shen, C., Wang, C. & Jia, W. Exosomes from beta-cells alleviated hyperglycemia and enhanced angiogenesis in islets of streptozotocin-induced diabetic mice. Diabetes Metab Syndr Obes 12, 2053–2064 (2019). 10.2147/DMSO.S213400

24 Mancini, A. D. et al. beta-Arrestin Recruitment and Biased Agonism at Free Fatty Acid Receptor 1. J Biol Chem 290, 21131–21140 (2015). 10.1074/jbc.M115.644450

25 Rakshit, K., Hsu, T. W. & Matveyenko, A. V. Bmal1 is required for beta cell compensatory expansion, survival and metabolic adaptation to diet-induced obesity in mice. Diabetologia 59, 734–743 (2016). 10.1007/s00125-015-3859-2

26 Sherman, B. T. et al. DAVID: a web server for functional enrichment analysis and functional annotation of gene lists (2021 update). Nucleic Acids Res 50, W216–W221 (2022). 10.1093/nar/gkac194

27 Szklarczyk, D. et al. STRING v10: protein-protein interaction networks, integrated over the tree of life. Nucleic Acids Res 43, D447–452 (2015). 10.1093/nar/gku1003

28 Shannon, P. et al. Cytoscape: a software environment for integrated models of biomolecular interaction networks. Genome Res 13, 2498–2504 (2003). 10.1101/gr.1239303

29 Subramanian, A. et al. Gene set enrichment analysis: a knowledge-based approach for interpreting genome-wide expression profiles. Proc Natl Acad Sci U S A 102, 15545–15550 (2005). 10.1073/pnas.0506580102

30 Javeed, N. et al. Pancreatic Cancer-Derived Exosomes Cause Paraneoplastic beta-cell Dysfunction. Clin Cancer Res 21, 1722–1733 (2015). 10.1158/1078-0432.CCR-14-2022

31 Midekessa, G. et al. Zeta Potential of Extracellular Vesicles: Toward Understanding the Attributes that Determine Colloidal Stability. ACS Omega 5, 16701–16710 (2020). 10.1021/acsomega.0c01582

32 Essandoh, K. et al. Blockade of exosome generation with GW4869 dampens the sepsis-induced inflammation and cardiac dysfunction. Biochim Biophys Acta 1852, 2362–2371 (2015). 10.1016/j.bbadis.2015.08.010

33 Kelpe, C. L. et al. Palmitate inhibition of insulin gene expression is mediated at the transcriptional level via ceramide synthesis. J Biol Chem 278, 30015–30021 (2003). 10.1074/jbc.M302548200

34 Manukyan, L., Ubhayasekera, S. J., Bergquist, J., Sargsyan, E. & Bergsten, P. Palmitate-induced impairments of beta-cell function are linked with generation of specific ceramide species via acylation of sphingosine. Endocrinology 156, 802–812 (2015). 10.1210/en.2014-1467

35 Xu, Y. N. et al. Low-grade elevation of palmitate and lipopolysaccharide synergistically induced beta-cell damage via inhibition of neutral ceramidase. Mol Cell Endocrinol 539, 111473 (2022). 10.1016/j.mce.2021.111473

36 Galadari, S., Rahman, A., Pallichankandy, S., Galadari, A. & Thayyullathil, F. Role of ceramide in diabetes mellitus: evidence and mechanisms. Lipids Health Dis 12, 98 (2013). 10.1186/1476-511X-12-98

37 Kakazu, E., Mauer, A. S., Yin, M. & Malhi, H. Hepatocytes release ceramide-enriched pro-inflammatory extracellular vesicles in an IRE1alpha-dependent manner. J Lipid Res 57, 233–245 (2016). 10.1194/jlr.M063412

38 Fukushima, M. et al. StAR-related lipid transfer domain 11 (STARD11)-mediated ceramide transport mediates extracellular vesicle biogenesis. J Biol Chem 293, 15277–15289 (2018). 10.1074/jbc.RA118.002587

39 Hirsova, P. et al. Lipid-Induced Signaling Causes Release of Inflammatory Extracellular Vesicles From Hepatocytes. Gastroenterology 150, 956–967 (2016). 10.1053/j.gastro.2015.12.037

40 Donoso-Quezada, J., Ayala-Mar, S. & Gonzalez-Valdez, J. The role of lipids in exosome biology and intercellular communication: Function, analytics and applications. Traffic 22, 204–220 (2021). 10.1111/tra.12803

41 Miyanishi, M. et al. Identification of Tim4 as a phosphatidylserine receptor. Nature 450, 435–439 (2007). 10.1038/nature06307

42 Record, M., Carayon, K., Poirot, M. & Silvente-Poirot, S. Exosomes as new vesicular lipid transporters involved in cell-cell communication and various pathophysiologies. Biochim Biophys Acta 1841, 108–120 (2014). 10.1016/j.bbalip.2013.10.004

43 Skotland, T., Sandvig, K. & Llorente, A. Lipids in exosomes: Current knowledge and the way forward. Prog Lipid Res 66, 30–41 (2017). 10.1016/j.plipres.2017.03.001

44 Kelleher, R. J., Jr., et al. Extracellular Vesicles Present in Human Ovarian Tumor Microenvironments Induce a Phosphatidylserine-Dependent Arrest in the T-cell Signaling Cascade. Cancer Immunol Res 3, 1269–1278 (2015). 10.1158/2326-6066.CIR-15-0086

45 Keller, S. et al. Systemic presence and tumor-growth promoting effect of ovarian carcinoma released exosomes. Cancer Lett 278, 73–81 (2009). 10.1016/j.canlet.2008.12.028

46 Lima, L. G., Chammas, R., Monteiro, R. Q., Moreira, M. E. & Barcinski, M. A. Tumor-derived microvesicles modulate the establishment of metastatic melanoma in a phosphatidylserine-dependent manner. Cancer Lett 283, 168–175 (2009). 10.1016/j.canlet.2009.03.041

47 Matsumura, S. et al. Subtypes of tumour cell-derived small extracellular vesicles having differently externalized phosphatidylserine. J Extracell Vesicles 8, 1579541 (2019). 10.1080/20013078.2019.1579541

48 Wieder, N. et al. FALCON systematically interrogates free fatty acid biology and identifies a novel mediator of lipotoxicity. Cell Metab 35, 887–905 e811 (2023). 10.1016/j.cmet.2023.03.018

49 Hastings, J. F., Skhinas, J. N., Fey, D., Croucher, D. R. & Cox, T. R. The extracellular matrix as a key regulator of intracellular signalling networks. Br J Pharmacol 176, 82–92 (2019). 10.1111/bph.14195

50 Musale, V., Wasserman, D. H. & Kang, L. Extracellular matrix remodelling in obesity and metabolic disorders. Life Metab 2 (2023). 10.1093/lifemeta/load021

51 Verrecchia, F. & Mauviel, A. Transforming growth factor-beta signaling through the Smad pathway: role in extracellular matrix gene expression and regulation. J Invest Dermatol 118, 211–215 (2002). 10.1046/j.1523-1747.2002.01641.x

52 Nomura, M. et al. SMAD2 disruption in mouse pancreatic beta cells leads to islet hyperplasia and impaired insulin secretion due to the attenuation of ATP-sensitive K+ channel activity. Diabetologia 57, 157–166 (2014). 10.1007/s00125-013-3062-2

53 Simeone, D. M. et al. Islet hypertrophy following pancreatic disruption of Smad4 signaling. Am J Physiol Endocrinol Metab 291, E1305–1316 (2006). 10.1152/ajpendo.00561.2005

54 Smart, N. G. et al. Conditional expression of Smad7 in pancreatic beta cells disrupts TGF-beta signaling and induces reversible diabetes mellitus. PLoS Biol 4, e39 (2006). 10.1371/journal.pbio.0040039

55 Lin, H. M. et al. Transforming growth factor-beta/Smad3 signaling regulates insulin gene transcription and pancreatic islet beta-cell function. J Biol Chem 284, 12246–12257 (2009). 10.1074/jbc.M805379200

56 Blum, B. et al. Reversal of beta cell de-differentiation by a small molecule inhibitor of the TGFbeta pathway. Elife 3, e02809 (2014). 10.7554/eLife.02809

57 Dhawan, S., Dirice, E., Kulkarni, R. N. & Bhushan, A. Inhibition of TGF-beta Signaling Promotes Human Pancreatic beta-Cell Replication. Diabetes 65, 1208–1218 (2016). 10.2337/db15-1331

