## Supplemental Material for "Lipotoxicity Induces Beta Cell Small Extracellular Vesicle-mediated β-cell Dysfunction"

**Running Title:** Lipotoxic β-cell EVs induce β-cell dysfunction

Abhishek Roy^1^, Alexandra Hoff^1^, Tracy K. Her^1^, Gallage Ariyaratne^1^, Kamalnath Sankaran Rajagopalan^1^, Matthew R. Brown, Alondra Soto-González, Aleksey V. Matveyenko^1,2^, Naureen Javeed^1,2*^

Supplemental Figures: 4

Supplemental Tables: 1

**Supplemental Methods**

**High fat diet studies.**

C57BL/6L mice were put on a high fat diet (HFD) or chow fed for 10 weeks. HFD islets were isolated and exposed to GW4869 (5 μM; S7609, Selleckchem) for 24 h (vs. untreated HFD islets). Islets were subjected to static glucose stimulated insulin secretion assays (Alpco) where islets were put in a 4 mM glucose solution made in Krebs Ringer buffer (KRB) for 30 minutes, then transferred to 16 mM glucose solution made in KRB for 30 minutes.

**TUNEL and Ki67 staining.**

C57BL/6L mouse islets were exposed to PAL EV (2X10^9^ every day for 48 h (vs. UT islets). Islets were then fixed in 4% paraformaldehyde. embedded in paraffin and sectioned. β-cell apoptosis and proliferation were assessed as previously conducted^4^ using a TUNEL In Situ CELL Death Detection kit (12156792910; Roche Diagnostics) and Ki67 antibody (550609; BD Pharmigen) along with insulin, respectively. Quantification of the percent TUNEL^+^ or Ki67^+^ β-cell/islet was analyzed using ImageJ.

**Supplementary Fig. 1**

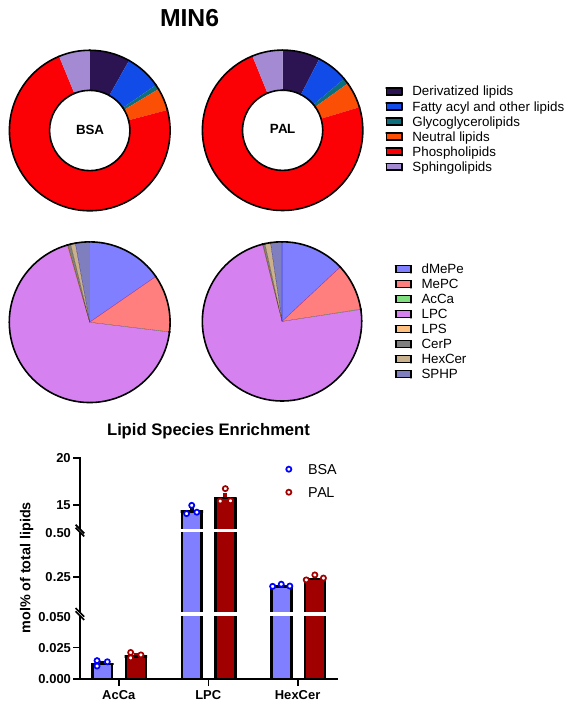

**Supplementary Fig. 1: Lipidomic analysis of palmitate stimulated MIN6 lysate.** MIN6 cells were treated with 0.5 mM palmitate or BSA (control) for 24 h. Lysates were subjected to lipidomic analysis (n=3 per condition). Differential expression of lipid species is defined as the molecular percentage of total lipids from BSA vs. PAL lysates. Values are a mean ± SEM. Statistical significance among groups is indicated by *, *p*<0.05.

**Supplementary Fig. 2**

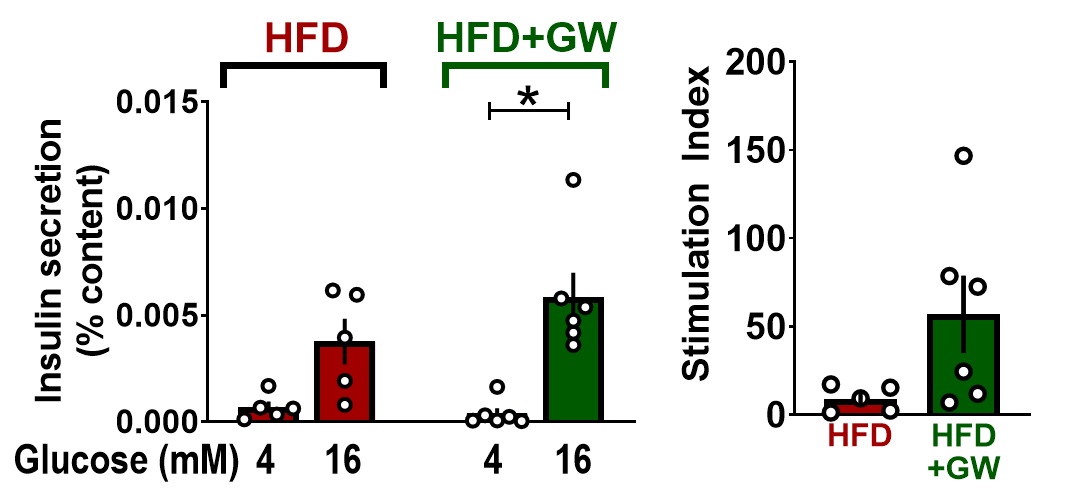

**Supplementary Fig. 2: Inhibition of sEV biogenesis in high fat diet islets improves GSIS.**  Isolated islets from C57BL/6L mice on a high fat diet (HFD) for 10 weeks were exposed to GW4869 (24h; vs. untreated islets). Static glucose stimulated insulin secretion (GSIS) was assessed at 4 mM basal and 16 mM stimulatory glucose concentrations and insulin stimulation index is expressed as 16 mM glucose divided by 4 mM basal concentrations (n=5-6 independent experiments per condition). Values are a mean ± SEM. Statistical significance among groups is indicated by *, *p*<0.05.

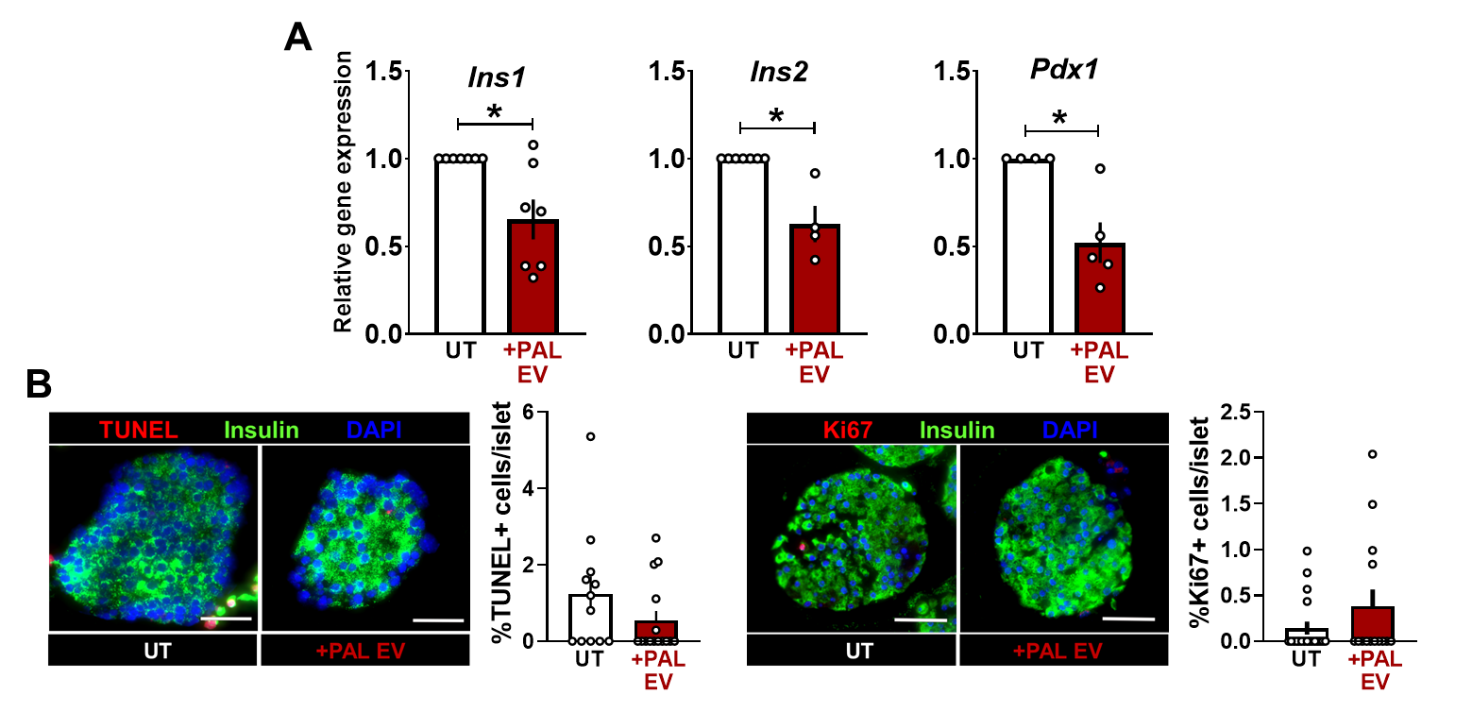
**Supplementary Fig. 3**

**Supplementary Fig. 3. Lipotoxic β-cell small EV do not induce β-cell apoptosis.** *A*: C57BL/6L mouse islets were exposed to PAL EV (2X10^9^ every day for 48 h (vs. UT islets) and assessed for gene expression of β-cell identity markers: *Ins1, Ins2*, and *Pdx1* (n=4-7 independent experiments). *B and C*: Paraffin-embedded C57BL/6L mouse islets treated with 2X10^9^ every day for 48 h (vs. UT islets) were sectioned and stained using immunofluorescence for TUNEL (apoptosis marker; red) or Ki67 (cell proliferation; red) and insulin (green). Quantification of β-cell apoptosis was assessed by TUNEL^+^ cells/islet in PAL EV treated islets vs. UT (*E*; n=12-15 total islets). Quantification of cell proliferation was measured by Ki67^+^ cells/islet (*F*; n=11-19 total islets). Values are a mean ± SEM. Statistical significance among groups is indicated by *, *p*<0.05.

**Supplementary Fig. 4**

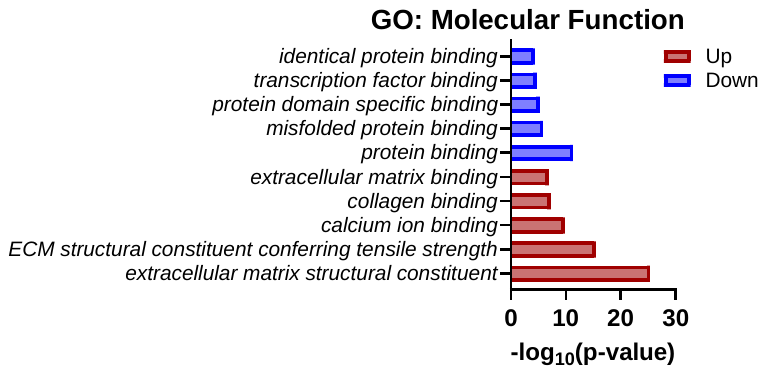

**Supplementary Fig. 4: Gene ontology (GO) analysis for Molecular Function.** C57BL/6L mouse islets were exposed to PAL EV (2X10^9^ EV/day; 48 h) vs. UT islets (n=2 biological repeats per condition). RNA-Sequencing was performed, then analyzed using Gene ontology (GO) looking at alterations in molecular function. PAL EV addition induced gene expression from pathways associated with ECM restructuring and binding (*p* < 0.05).

**Supplementary Table 1. Primers**

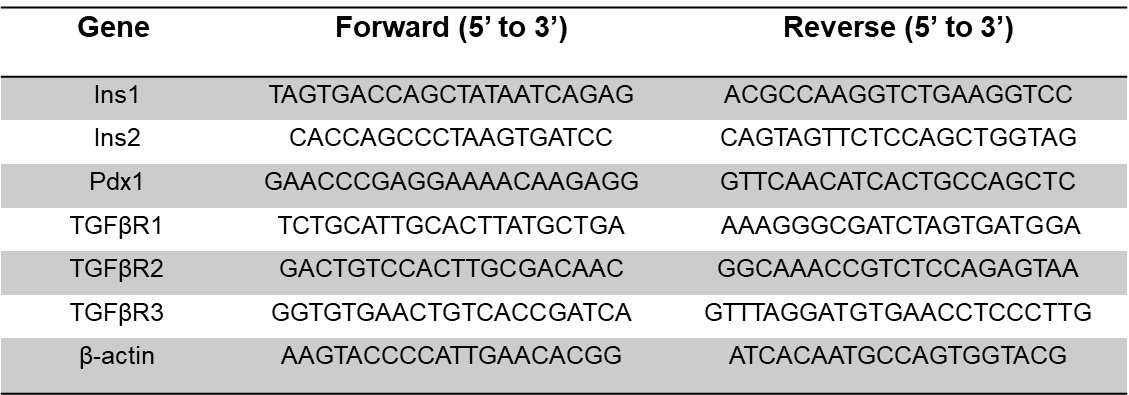
